# Genetic variation at the species and population levels in the Rocky Mountain ridged mussel (*Gonidea angulata*)

**DOI:** 10.1101/2020.11.16.385195

**Authors:** Karen E. Mock, James A. Walton, Steven F. R. Brownlee, Jon H. Mageroy, Greg Wilson, Ian R. Walker

## Abstract

Freshwater mussels in western North America are threatened by water diversions, climate change, loss of required host fish, and other factors, and have experienced marked decline in the past several decades. All four of the primary lineages (potentially species) of freshwater mussels in the western U.S. and Canada are widespread and have somewhat generalist host fish requirements. Of these lineages, perhaps the most poorly understood and of greatest conservation concern is *Gonidea angulata* (Rocky Mountain ridged mussel). *Gonidea* is a monotypic genus occurring only in the western continental U.S. and southern Canada. Here we describe the patterns of genetic variation across the species range, including several populations in the Okanagan Valley at the northern edge of the range. We detected only ten mitochondrial cytochrome oxidase I haplotypes, three of which ere commonly found across major hydrologic basins, and the remainder of which were basin-specific variants. Haplotypes differed by a maximum of 5 of 537 nucleotides. New microsatellite loci were developed for *G. angulata* as a part of this study. Data from these microsatellite loci indicated that the population in the Chehalis River, Washington, was distinct from other locations, and that the Okanagan lake population was somewhat diverged from the remaining populations in the Columbia River and Klamath Lake. Only low levels of inbreeding were detected, in contrast to previous findings in *Margartifera falcata,* suggesting that hermaphroditism is not common. The population with the least diversity, according to microsatellite data, was the northernmost known population in Okanagan Lake We discuss the biogeographic and conservation implications of our findings.

## Introduction

Freshwater mussels (Unionida) provide a range of critical ecosystem services related to filtration (e.g. affecting water clarity, productivity, sedimentation rates, nutrient and energy transfer to benthic ecosystems) and substrate structural diversity (e.g. providing stabilization and enhancing biodiversity) (Lopes-Lima et al. 2014, Vaughn 2017). Freshwater mussels also have a range of cultural values, both historically and currently (Vaughn 2017). However, these animals are declining worldwide at an alarming rate (Bogan 2008). North America is the global center of species diversity for freshwater mussels, but also the center of their decline (Williams et al. 1993; Strayer et al. 2004). Freshwater mussels rely on fish for larval development and dispersal, and many have highly specialized relationships with a narrow range of host fish species. This reliance on fish hosts compounds their vulnerability. Further, as sessile filter feeders, freshwater mussels are susceptible to sedimentation, depleted oxygen levels, temperature increases, and pollutants, all common freshwater impacts from human activities.

While the southeastern US is the global epicenter of freshwater mussel species diversity, the western US and Canada also contain freshwater mussel species which were historically widespread and important both ecologically and culturally (Brim Box et al. 2006; Brubaker et al. 2009). These species are also experiencing rapid rangewide declines of unknown etiology across their ranges, and are increasingly imperiled (Blevins et al. 2016a, 2016b, 2016c, 2016d, Vinarski and Cordeiro 2011).

There are currently four major lineages of freshwater mussels in western Canada and the western contiguous US: (1) a lineage consisting of *Anodonta nuttalliana* (Lea, 1838) (winged floater)*, A. californiensis* (I. Lea, 1852) (California floater)*, and A. wahlamatensis* (I. Lea 1938), here referred to as *A. nuttalliana* (Chong et al. 2008), (2) a lineage consisting of *Anodonta oregonensis* (I. Lea, 1838) (Oregon floater) and *A. kennerlyi* (I. Lea, 1860) (western floater), here referred to as *A. oregonensis* (Chong et al. 2008), (3) *Margaritifera falcata* (Gould 1850) (western pearlshell), and (4) *Gonidea angulata* (I. Lea 1838) (Rocky Mountain ridged mussel or western ridged mussel). In western North America, rangewide genetic patterns in the two most broadly distributed species, *A. nuttalliana* and *M. falcata*, have been reported and compared. Rangewide genetic patterns in *A. nuttalliana* revealed pronounced genetic structure consistent with long-term subdivision in major hydrologic basins of the western U.S. (Mock et al. 2010). There was also a lack of support for morphologically-based subdivisions which had been the basis of taxonomy in this group.

By contrast, rangewide genetic patterns in *M. falcata* showed very little geographic structuring in genetic diversity but pronounced levels of inbreeding in many populations (Mock et al. 2013). The signature of inbreeding may be due to hermaphroditism in this species (Kat 1983), and is expected to increase as populations become more isolated. Because salmonid fish are a major host for *M. falcata*, declines in the numbers and dispersal distances of salmonids may be contributing to isolation of *M. falcata* populations. Conservation recommendations based on this work include (i) placing less restriction on transfers of *M. falcata* across hydrologic basins, but (ii) assessing source population diversity prior to translocations, (iii) increasing genetic diversity in isolated populations through translocation, and (iv) assessing host fish numbers and dispersal patterns (e.g. fish barriers) in streams where *M. falcata* is present.

*G. angulata* is also found only in western North America, and has several anatomical characters that are unique among North American freshwater mussel fauna (Ortmann 1916). *G. angulata* is the least well-studied of the freshwater mussels in western Canada and the contiguous western U.S., but is perhaps the species of greatest conservation concern in those landscapes, as *Gonidea* is a monotypic genus, and this single species represents a considerable portion of the evolutionary diversity within the Family Unionidae (Lopes-Lima et al. 2017). Our goal in this study was to provide an assessment of rangewide genetic diversity and divergence in *G. angulata*, with attention to both geographic structuring and inbreeding patterns, and how they relate to life history traits, hydrologic basin geography, and patterns in other western freshwater mussels.

## Study System

*G. angulata* is historically thought to have occurred from central California northward across Oregon and Washington, into southern British Columbia and east into northern Nevada, Idaho, and possibly into northwestern Montana (Gangloff and Gustafson 2000) and northwestern Utah (Henderson 1924, 1929, 1936). Dramatic declines have been reported in California (Howard et al. 2015), and throughout the Columbia and Snake River basins (Brim Box et al. 2006, Frest and Johannes 1995) as well as in Nevada and Idaho. The species is now thought to be extirpated from central California, Montana, and Utah (NatureServe Explorer, accessed 25 November 2019). *G. angulata* is listed at state/province, national, and international levels as a species of concern (Table 1). In Canada, *G. angulata* is restricted to the Okanagan Valley (COSEWIC 2010).

**Table 1.** Current conservation status of *Gonidea angulata.*

| Level | Status |
| --- | --- |
| Global | G3: Vulnerable throughout range (NatureServe 2007) |
| Global | Vulnerable (International Union for Conservation of Nature (IUCN) Red List of Threatened Species 2017) |
| Canada | N2: Nationally Imperiled (NatureServe 2017)<br>www.natureserve.org/conservation-tools/conservation-status-assessment |
| Canada | Endangered (Committee on the Status of Endangered Wildlife in Canada (COSEWIC) 2010) |
| Canada | SC: Species of Special Concern (Canadian Species at Risk Act (SARA) 2005)<br>Proposed for reclassification to Endangered (SARA 2019)<br>www.gazette.gc.ca/rp-pr/p1/2019/2019-03-23/html/reg1-eng.html |
| USA | N3: nationally Vulnerable (NatureServe 2004) |
| British Columbia | S2: Imperiled (NatureServe 2014)<br>British Columbia Species and Ecosystems Explorer 2014)<br>www2.gov.bc.ca/gov/content/environment/plants-animals-ecosystems/conservation-data-centre/explore-cdc-data/species-and-ecosystems-explorer |
| Washington | S2S3: Vulnerable in Washington (NatureServe 2009)<br>SGCN: Species of Greatest Conservation Need (WDFW 2015)<br>Sensitive (USFS Region 6 2019)<br>Sensitive (BLM 2019) |
| Oregon | S2S3: Imperiled to Vulnerable in Oregon (NatureServe)<br>CS: Conservation Strategy Species (ODFW 2016)<br>List 1: Threatened or Endangered Throughout Range (ORBIC 2016)<br>Sensitive (USFS Region 6 2019)<br>Sensitive (BLM 2019) |
| Idaho | S3: Vulnerable in Idaho (NatureServe 2015) |
| Nevada | S1: Vulnerable in Nevada (NatureServe 2019)<br>Sensitive (BLM 2019) |
| California | S1S2: Critically Imperiled to Imperiled in California (NatureServe 2005) |

*G. angulata* is found in streams, rivers, and lakes, in a broad range of substrate types (Taylor 1981; Frest and Johannes 1995), but seems to require cold, clear, oligotrophic waters with constant flow (COSEWIC 2003, 2010). Individuals of the species can be long-lived; perhaps 20-30 years or more (Vannote and Minshall 1982). As with other unionid mussels, *G. angulata* requires a host fish for development of glochidia and dispersal to new habitats. Sculpin (*Cottus* spp.) are likely the primary host fish for *G. angulata,* although several other species may serve as secondary hosts in local areas (COSEWIC 2010; Stanton et al. 2012; O’Brien et al. 2013, Mageroy 2015a). *Cottus* species are distributed across the northern hemisphere, including throughout the western U.S. and Canada (Froese and Pauly 2019), and frequently co-occur with salmonid species, requiring relatively clean, cold water, and an extensive phylogeographic study of *Cottus* species is currently underway (https://www.fs.fed.us/rm/boise/AWAE/projects.html). Freshwater *Cottus* species are generally small, benthic fish, and unlikely to serve as individual long-distance dispersers of glochidia, although other host fish may provide glochidial dispersal (Martel and Lauzon-Guay 2005). Currently, only limited information about host fish suitability exists for *G. angulata*, but Pit roach (*Lavinia symmetricus mitrulus*), longnose dace (*Rhinichthys cataracta*), leopard dace (*Rhinichthys falcatus*), and northern pikeminnow (*Ptychocheilus oregonensis*), hardhead (*Mylopharodon conocephalus*), and Tule perch (*Hysterocarpus traski*) have been implicated as secondary hosts in some local studies (Haley et al. 2007; Spring Rivers 2007; Stanton et al. 2012; Magerøy 2015a). Additionally, occasional hermaphroditism has been noted in *G. angulata* (van der Schalie 1970), suggesting that in some isolated populations, inbreeding could be a concern, as it is for *M. falcata* (Mock et al. 2013).

## Methods

### Sample Collection and DNA Extraction

We sampled 270 *G. angulata* samples (foot clips), collected non-lethally from five major river basins across the species’ range in the western United States: Columbia (Washington, Oregon, and Idaho; n=98), Chehalis (western Washington; n=21), Klamath (western Oregon; n=55), Sacramento (central California; n=5) and Humboldt (Nevada; n=5) (Figure 1a, Table S1). Our samples also included foot clips from 6 locations in the Okanagan River Basin (a Columbia River tributary; n=86) (Figure 1b, Table S1) in British Columbia, Canada, to assess finer-scale genetic variation. The Columbia, Chehalis, Klamath and Sacramento Rivers all flow westward to the Pacific Ocean. The Humboldt River is part of a closed basin system, draining ultimately into the Humboldt Sink of northeastern Nevada. In Canada, *G. angulata* are known only from the Okanagan Valley; more specifically from the Okanagan River and the four lakes (Okanagan Lake, Skaha Lake, Vaseux Lake, and Osoyoos Lake) connected by this river (Figure 1b). These lakes vary greatly in size, ranging from Okanagan Lake (approximately 100 km long, and 273 m deep) to Vaseux Lake (3.8 km long, and 27 m deep). All samples were preserved in 95% ethanol.

**Figure 1.**
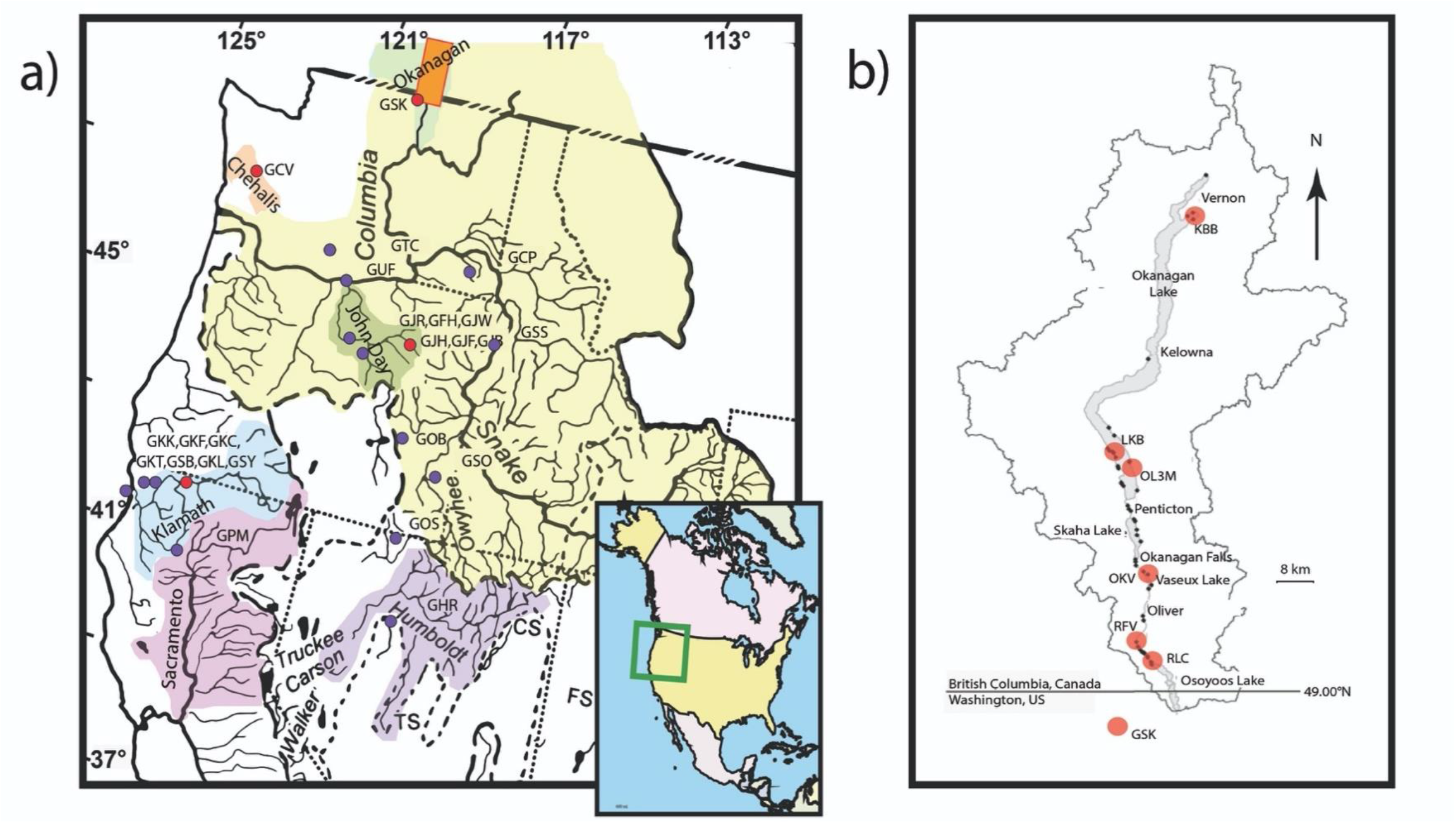
Maps showing approximate location of sampling sites and distribution of Rocky Mountain ridged mussel in the United States and Canada. a) Western United States: colored regions indicate major watersheds and sub-basins with known occurrences of the mussel. Orange rectangle indicates Okanagan region in Canada, purple dots indicate US genetic sampling locations with COI data, red dots indicate sites with both μsat and COI data. Green square on inset map of North America (lower right) indicates approximate bounds of the larger map in North America; b) Okanagan Watershed in Canada. Red dots indicate Canadian genetic sampling locations (all sites have both μsat and COI data), black dots indicate known mussel populations.

### Mitochondrial Sequencing

Genomic DNA was extracted from each of the subsampled tissues using a Qiagen DNeasy Blood and Tissue kit following the manufacturer protocol, with negative controls in each separate extraction group. We obtained mitochondrial female-lineage cytochrome c oxidase I subunit (COI) sequences, approximately 650 base pairs (bp) in length, from 3-10 samples in each population. Amplification was performed using the HCO700dy (Hoeh et al. 2002; 5’ TCAGGGTGACCAAAAAATCA, HCO2198 first 5’ end 6 bases removed) and LCO1490 (Folmer et al. 1994; 5’ GGTCAACAAATCATAAAGATATTGG) primers. Amplification reactions contained 1x MyTaq HS Master Mix (Bioline™), 0.1 mg/mL BSA, 0.5 μM of each forward and reverse primer, and approximately 15 ng genomic DNA. Reactions were denatured at 95 °C for 2 min., followed by 35 cycles of 94 °C for 30 s, 50 °C for 60 s, 72 °C for 90 s, with a final extension step of 72 °C for 5 minutes. Reactions were then analyzed for quality assurance via 1.4 % agarose gel prior to sequencing. Bidirectional sequences were obtained using primer pairs HCO700dy and internal primer LCO1550 or LCO1490 (Chong et al. 2008). Contiguous COI sequences, using forward and reverse sequencing primers, were constructed using Geneious Pro v.5.6.7 (https://www.geneious.com) for alignment, editing, and quality assessment. Contiguous sequences for each individual were aligned using MEGA4 software (Tamura et al. 2007) and trimmed to 537 bp. Genealogical relationships among mitochondrial COI haplotypes were estimated using TCS v1.21 (Clement et al. 2000).

### Microsatellite Development and Optimization

Microsatellites, are regions of the nuclear genome that are hypervariable and biparentally inherited, so they are excellent genetic loci for assessing population divergence, gene flow, diversity, and inbreeding. These loci require initial development from ‘shotgun’ genomic sequences, followed by refinement, primer design, and population-level assessment. Our methods for development are described in detail in Documents S1-S5. A final set of 18 polymorphic microsatellite loci developed and optimized for multiplexing (Table S2). These include 10 loci with dimeric repeats, 4 with trimeric repeats, and 4 with tetrameric repeats.

### Population Genetic Analysis

We used microsatellite genotypes to assess genetic divergence among populations, genetic diversity within populations, and the strength of assignment of individuals to populations. Multilocus microsatellite genotypes (Document S6) were obtained from 169 samples representing three major hydrologic basins (Columbia, Chehalis, Klamath) and 10 populations (Table S1).

For the eight populations with at least 16 individuals represented, Hardy-Weinberg equilibrium (HWE) was assessed using the exact test with a Markov chain method (Guo and Thompson, 1992) in GenePop v4.7.0 (Raymond and Rousset 1995; Rousset 2008). HWE was also assessed with heterozygote deficiency as a specific alternative hypothesis (Rousset and Raymond 1995). Null allele frequencies were estimated using a maximum likelihood estimation (Dempster et al. 1977) as implemented in GenePop v4.7.0. Diversity indices Na (number of alleles observed across all loci), Ne (number of effective alleles), Ho (observed heterozygosity), uHe (unbiased estimate of expected heterozygosity if genotypes are in Hardy-Weinberg proportions), and F (fixation index) were estimated using GenAlEx software (Peakall and Smouse 2006, 2012).

Allelic richness (AR), which can be influenced by sample size, was estimated using a rarefaction approach using FSTAT v2.9.4 (Goudet 2003). Separate estimates of AR were made using minimum population sizes of either 5 or 13 individuals per population with complete genotypes. The RFV population contained only 5 genotypes with no missing data. Genetic differentiation across and between all populations, *Θ*_ST_ was estimated using FSTAT v2.9.4 (Goudet 2003), bootstrapping over loci to obtain 95% confidence intervals. An individual-based population assignment/cross-assignment test was performed using GenAlEx software (Peakall and Smouse 2006, 2012), with the “leave-one-out” option, to characterize patterns of similarity among populations. Only samples with complete genotypes were used for assignment analysis.

We assessed the signature of recent bottlenecks in all populations except RFV and RLC, which had sample sizes too low for this kind of inference. A recent bottleneck is expected to reduce allelic richness more rapidly than heterozygosity, resulting in an excess of heterozygosity relative to expectations for the observed number of alleles under mutation-drift equilibrium (Cornuet and Luikart 1997). Furthermore, a bottleneck is expected to produce a shift in the frequency distribution of alleles (Luikart et al. 1998). We used Bottleneck v1.2.02 (Cornuet and Luikart 1997) to determine whether either of these signatures was present in populations, applying the sign test, a standard differences test, a Wilcoxan 1-tailed test for heterozygosity excess, and a mode-shift test, assuming a two-phase mutational model, and using a coalescent simulation.

Individual-based population structuring using a Bayesian approach was also performed using the program Structure v2.3.4 (Pritchard et al. 2000) using the genotype dataset with no missing data (Document S6). A burn-in period of 20,000 MCMC iterations was followed by 10,000 iterations in the Structure program, with a uniform prior for the alpha parameter, assuming allele frequency correlation among populations, and using no prior location information. This number of iterations was sufficient for parameter estimates to stabilize in the model. Each value of k (number of clusters) from 1-10 was assessed, with 10 replications per k. The optimal level of k was identified using the delta-K method of Evanno et al. (2005), implemented in Structure Harvester (Earl and vonHoldt 2012).

## Results

### Mitochondrial Sequencing

A total of 152 bidirectional COI sequences (Tables 2, S3) were obtained from a total of 30 sampling sites, including 36 from Okanagan River Basin populations, 61 from an array of Columbia River tributary populations in Washington, Idaho, and Oregon, 40 from the Klamath River Basin, and five each from the Humboldt and Sacramento Basins. A total of ten different haplotypes were detected across our dataset, none separated by more than 5 (~0.8%) mutational differences over the 537 bp alignment (Figure 2). Three geographically widespread haplotypes were detected (GonA, GonB, and GonD). GonA was the most widespread and putatively ancestral haplotype, occurring in all five major basins (Figure 2). GonB was found in the Humboldt river population, and was widely distributed in the Columbia watershed, but was not detected in the Chehalis, Klamath or Sacramento populations. The Klamath River populations contained a very common and unique haplotype (GonH; 30/39 samples) and was the most divergent of the sampling sites. The Chehalis River also contained a unique haplotype (GonF) which was present in three of the five samples from that population, suggesting a distinct haplotype frequency. Other haplotypes unique to populations were found only in low numbers, which are vulnerable to sample size artifacts. Although sample numbers were similar across sites, there were four sites in which we detected only a single haplotype (KBB in the Okanagan River Basin, GKT and GSB in the Klamath Basin, GPM in the Pit River).

**Figure 2.**
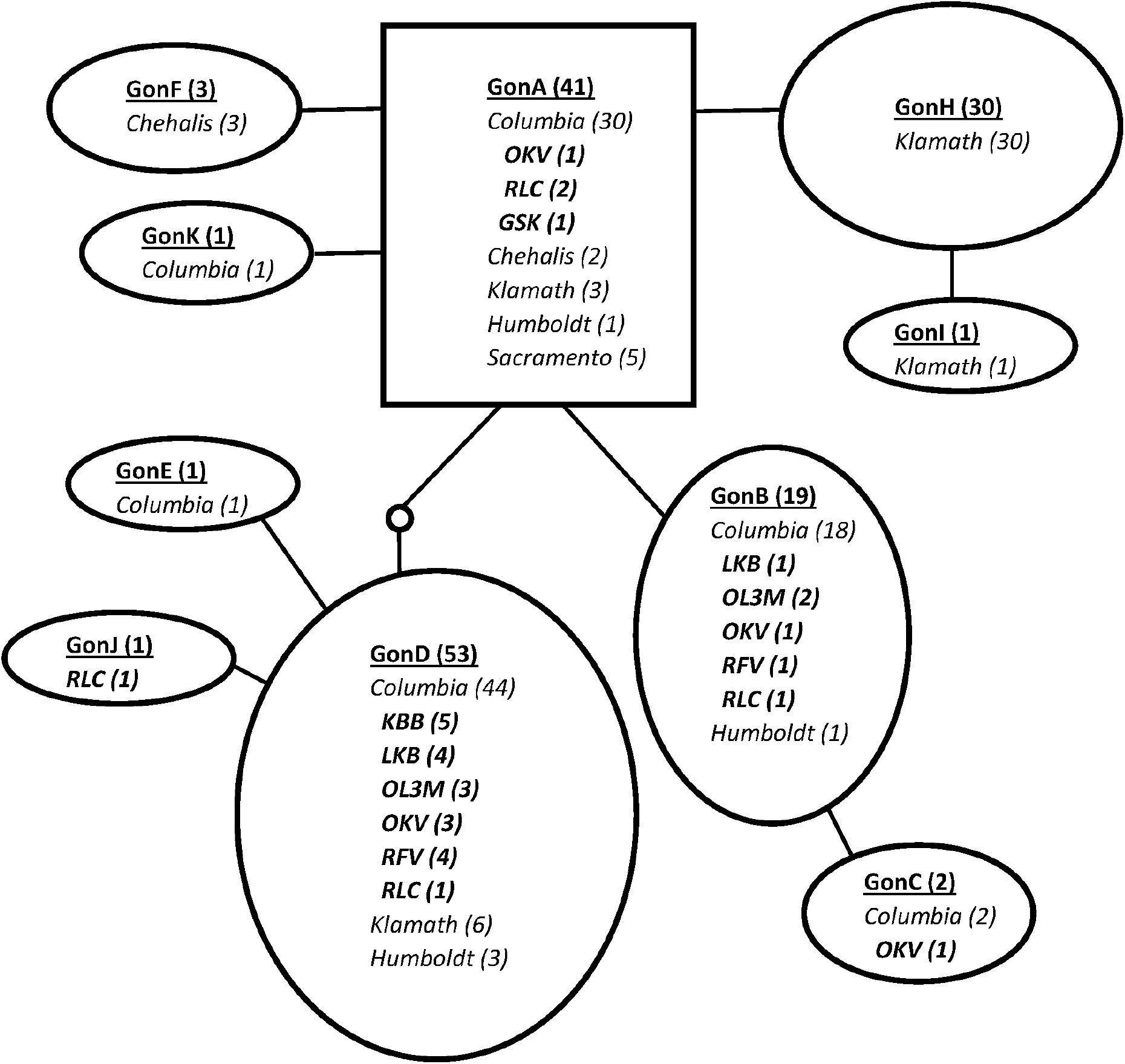
Network of mitochondrial COI haplotypes observed among sampling locations. Okanagan Basin populations are designated in bold italics. The rectangle represents the likely ancestral haplotype. Lines represent single mutational differences among haplotypes. The small circle represents an unobserved haplotype. Numbers in parentheses represent the numbers of haplotypes observed by location. The figure was created using TCS software (Clement et al. 2000).

**Table 2.**
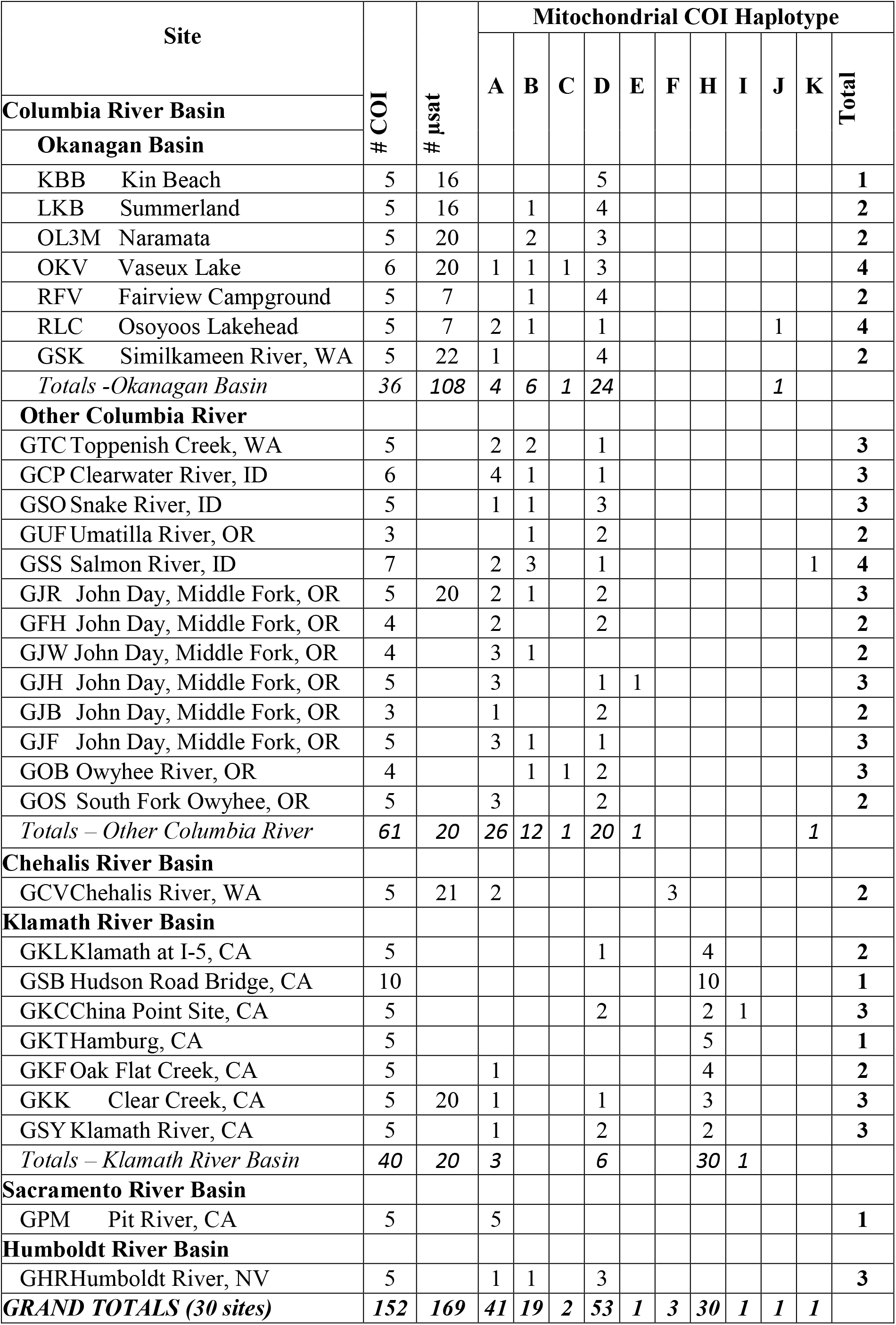
Distribution of mitochondrial cytochrome oxidase I (COI) haplotypes across sites and total number of COI sequences and microsatellite (μsat) genotypes by population.

The haplotype GonD was particularly common in, but not unique to, the Okanagan River Basin populations, occurring in every sampled site. GonD was the only haplotype detected at Kin Beach (KBB). The GonD haplotype was also detected in 3 of the 5 samples from the Humboldt Basin (GHR). A haplotype unique (GonJ) to the Okanagan Valley was detected in a single individual at the Lakehead Campsite (RLC) (Table 2).

### Population Genetic Analyses

No identical genotypes were detected within or across populations, and individual genotypes differed by a minimum of 16 allelic mismatches. Null allele frequency estimates by population ranged from 0.04 to 0.07, with the highest rates occurring in the GAN8467 locus, where the null allele frequency averaged 0.23 across populations. Null alleles in all other loci ranged from 0.01 to 0.12. We did not consider these rates to be high enough to warrant locus exclusion.

Pronounced HWE deviations were noted in populations KBB and OKV, relative to the random mating model (Table 3). When heterozygote deficiency was the alternative hypothesis in the HWE assessment, all populations except GCV showed significant deviations from HWE expectations under random mating, suggesting that heterozygote deficiencies are widespread (Table 3). All populations showed somewhat reduced heterozygosity relative to expectations (Table 4). Allelic richness was similar among populations, although GCV and KBB were consistently low. With rarefaction to n=5, RFV, RLC, and GJR had the greatest AR, but the power to detect differences would be low at this level. With rarefaction to n=13 (excluding RFV and RLC), GJR, OKV, GSK, and GKK had the highest AR. Population-level inbreeding (estimated by F, the fixation index, Table 4) was low but positive for all populations, and above 0.10 in KBB and OKV.

**Table 3.** Hardy-Weinberg equilibrium assessment for *G. angulata* populations represented by >16 samples. Probability estimates of observed data given random mating (rm), and with the alternative hypotheses of heterozygote deficiency.

| Population | Prob.<br>$H_0 =$<br>rm | Prob.<br>$H_0 =$ rm<br>$H_1 =$ het. deficit |
| --- | --- | --- |
| KBB | 0.000 | 0.000 |
| LKB | 0.102 | 0.002 |
| OLM3 | 0.069 | 0.000 |
| OKV | 0.001 | 0.000 |
| GCV | 0.152 | 0.479 |
| GSK | 0.021 | 0.000 |
| GJR | 0.088 | 0.000 |
| GKK | 0.179 | 0.002 |

**Table 4.** Diversity metrics by population in *G. angulata*, based on microsatellite data: N (number of sampled individuals); Year (year collected); Na (number of alleles observed across all loci); Ne (number of effective alleles); AR (average allelic richness across loci, estimated using rarefaction with a minimum population size of 5 or 13 individuals with complete data); Ho (observed heterozygosity); uHe (unbiased estimate of expected heterozygosity if genotypes are in Hardy-Weinberg proportions); F (fixation index).

| Pop | Year | N | Na (SE) | Ne (SE) | AR5 | AR13 | Ho (SE) | uHe (SE) | F (SE) |
| --- | --- | --- | --- | --- | --- | --- | --- | --- | --- |
| KBB | 2016 | 16 | 4.8 (0.2) | 3.1 (0.2) | 3.73 | 4.70 | 0.57 (0.04) | 0.67 (0.02) | 0.13 (0.06) |
| LKB | 2016 | 16 | 7.0 (0.5) | 4.3 (0.4) | 4.68 | 6.67 | 0.67 (0.04) | 0.75 (0.03) | 0.08 (0.04) |
| OL3M | 2017 | 20 | 7.5 (0.6) | 4.6 (0.4) | 4.74 | 6.69 | 0.69 (0.04) | 0.77 (0.02) | 0.08 (0.04) |
| OKV | 2016 | 20 | 9.8 (0.6) | 6.1 (0.5) | 5.62 | 8.56 | 0.72 (0.04) | 0.83 (0.02) | 0.11 (0.04) |
| RFV | 2017 | 7 | 6.8 (0.4) | 5.2 (0.4) | 5.77 | na | 0.76 (0.04) | 0.85 (0.02) | 0.04 (0.05) |
| RLC | 2016 | 7 | 6.7 (0.3) | 5.2 (0.4) | 5.75 | na | 0.76 (0.05) | 0.85 (0.02) | 0.04 (0.05) |
| GSK | 2006 | 22 | 9.9 (0.6) | 5.6 (0.5) | 5.46 | 8.38 | 0.72 (0.04) | 0.81 (0.02) | 0.09 (0.05) |
| GCV | 2007 | 21 | 6.2 (0.7) | 3.9 (0.5) | 3.98 | 5.47 | 0.67 (0.05) | 0.70 (0.03) | 0.02 (0.05) |
| GJR | 2007 | 20 | 10.2 (0.7) | 6.4 (0.6) | 5.73 | 8.88 | 0.72 (0.04) | 0.83 (0.02) | 0.10 (0.05) |
| GKK | 2008 | 20 | 9.3 (0.6) | 5.3 (0.6) | 5.28 | 8.11 | 0.72 (0.04) | 0.78 (0.03) | 0.05 (0.02) |

We did not detect the signature of a recent population bottleneck in any of the sampled populations except KBB, the northernmost extremity of the species distribution. Okanagan River basin populations, which showed a low probability of being in mutation-drift equilibrium using the standard differences test (p = 0.03), but no deviations using the sign test, the Wilcoxan 1-tailed test, or the mode shift test.

Structuring among populations was low but detectable, with an overall *Θ*_ST_ of 0.083 (95% c.i. 0.065-0.104). GCV, from the Chehalis River Basin, was the most distinct population, showing clear differentiation in the PCoA and Structure analyses (Figures 3,4). The Okanagan Lake populations (KBB, LKB, OL3M) were similar to each other relative to the other Okanagan River Basin populations, but there was evidence of mixing between Okanagan Lake and downstream populations OKV, RFV, and RLC (Figures 3,5). GKK, the Klamath population, shows affiliation with the Columbia River Basin populations GJR and GSK, which are geographically much closer to the Okanagan River Basin population than GKK (Figures 1,3,5), but this affiliation was not strong enough to be reflected in the population assignment results (Table 5). When assessed separately, the Okanagan River Basin populations showed little structuring, although the KBB population was distinct (Figure 3). Individual-based assignment tests showed that individuals from the KBB, GCV, and GKK populations showed 100% self-assignment, suggesting restricted gene flow, and across populations the assignment test results were generally consistent with the PCoA results.

**Figure 3.**
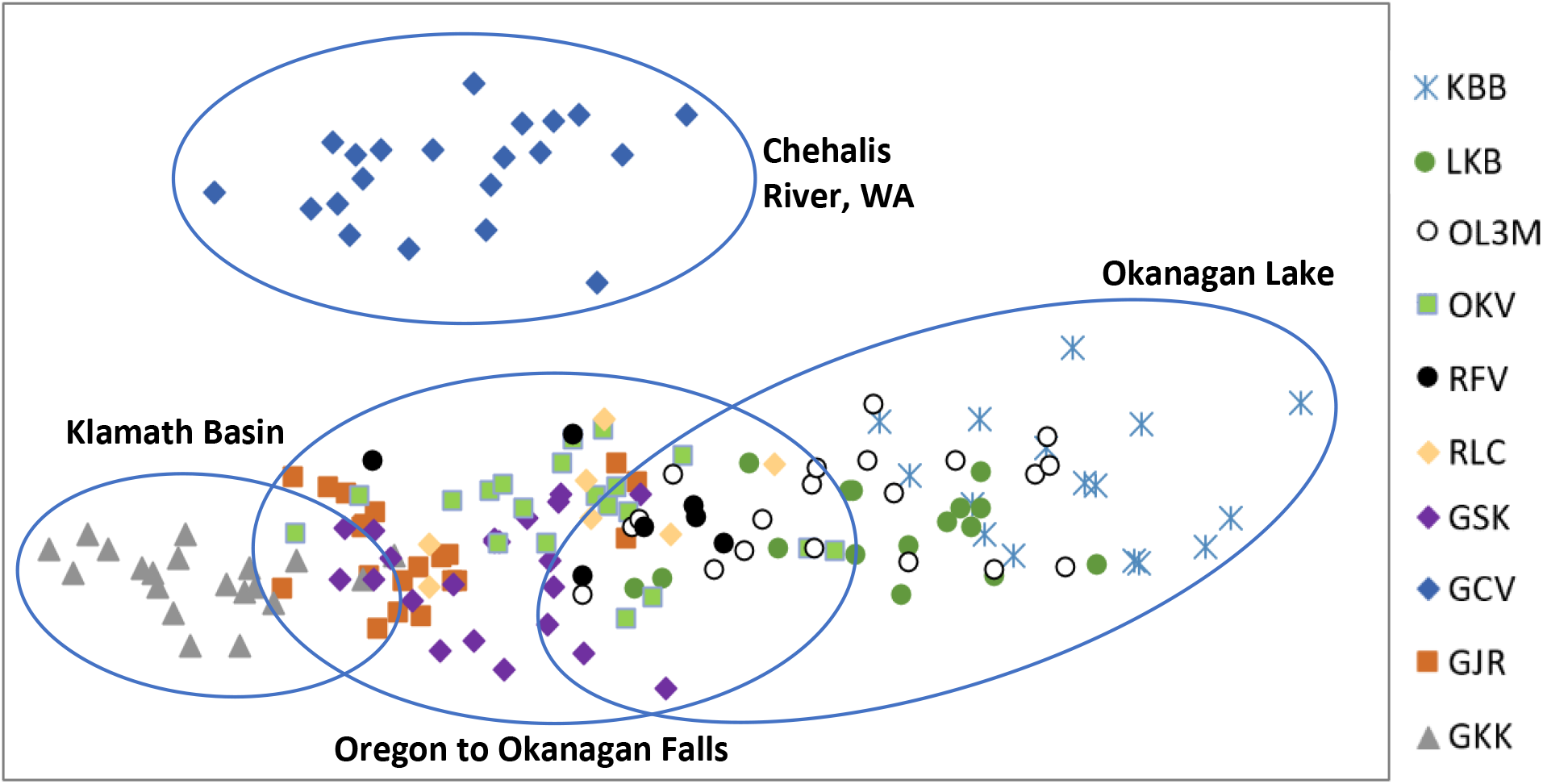
Principal coordinates analysis of individuals in all study populations based on nuclear microsatellite loci. The first two axes explained 11.15% of the total variation in the dataset.

**Figure 4.**
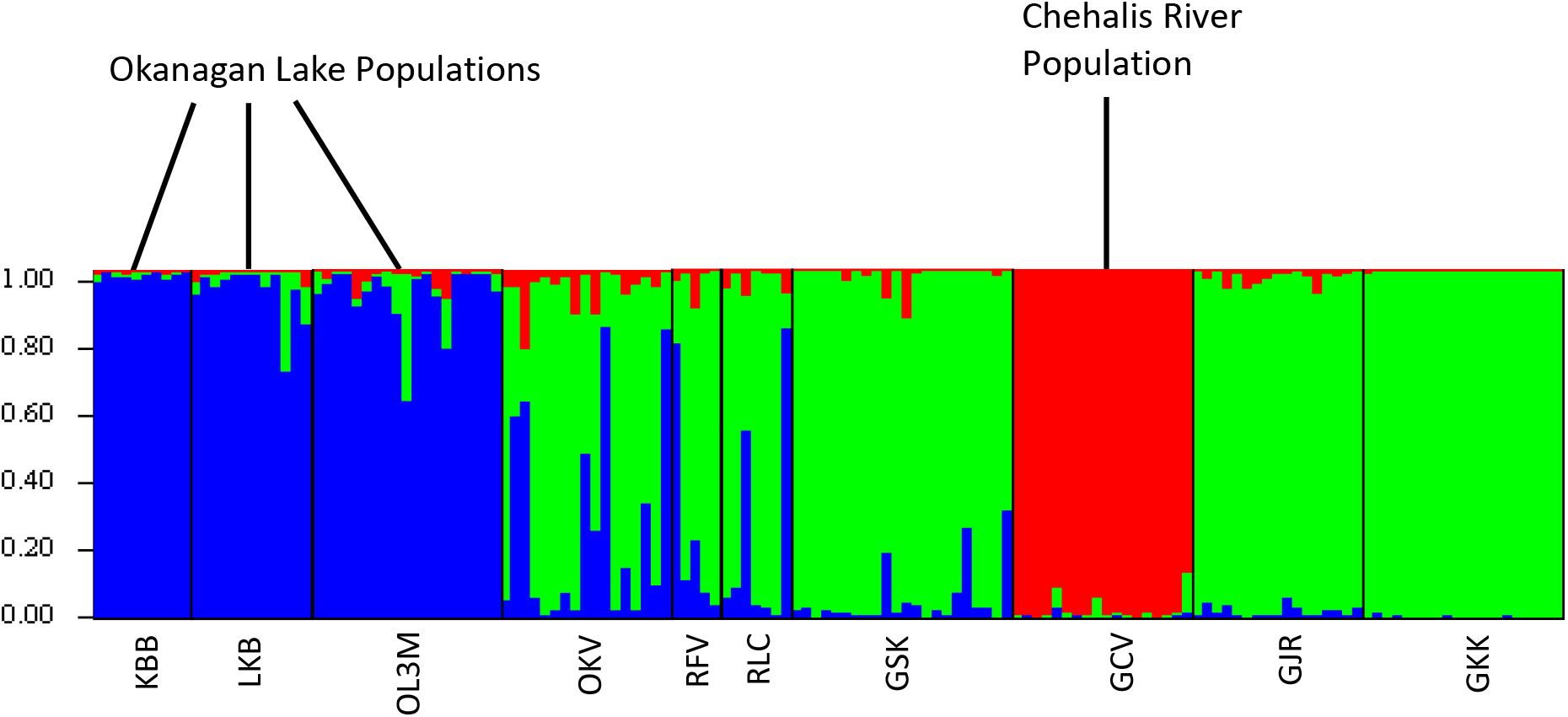
Bar plot of STRUCTURE results for the k=3 run with the highest posterior probability. Each vertical bar represents an individual, and the proportion of each of the three colors in each bar represents the probability of membership in each of the three hypothesized groups. Vertical black lines bound groups of individuals in each sampled population, and sample population abbreviations are below these groups.

**Figure 5.**
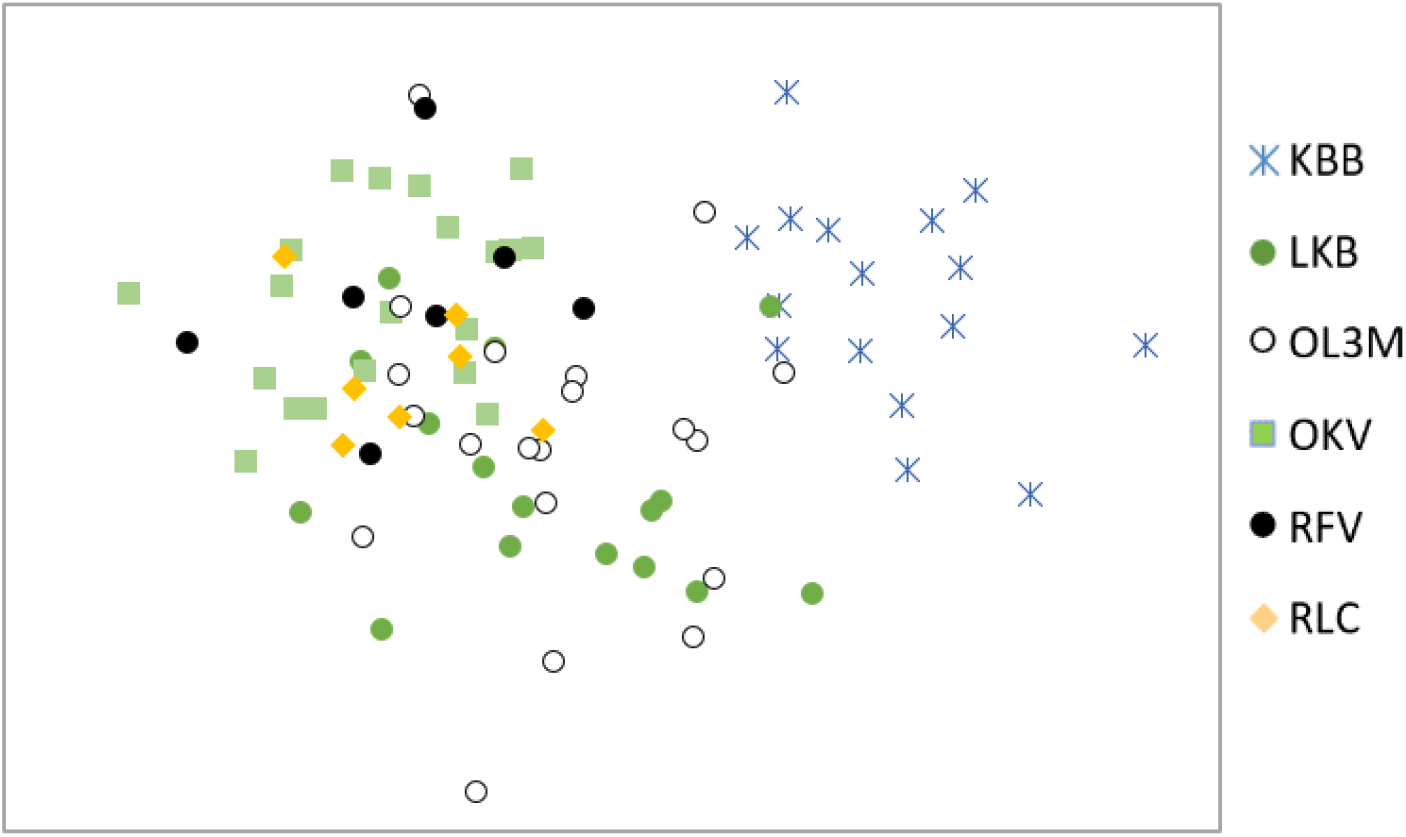
Principal coordinates analysis of individuals from Okanagan Basin populations in Canada, based on nuclear microsatellite analysis. The first two axes explained 11.62% of the variation in the dataset.

**Table 5.** Individual-based population assignment results based on *G. angulata* microsatellite genotypes. Populations marked with an asterisk had low sample size numbers, reducing the probability of accurate self-assignment. Only individuals with no missing allelic data were used in this analysis.

| Population | Self-assignment | Assignment to other populations |
| --- | --- | --- |
| KBB | 10 | 0 |
| LKB | 6 | OL3M(6) |
| OL3M | 15 | LKB(4) |
| OKV | 8 | RFV(4), LKB(1), OL3M(1), RLC(2), GSK(1) GJR(1) |
| RFV* | 0 | OKV(3), RLC(1), LKB(1) |
| RLC* | 0 | OKV(3), RFV(1), GSK(1), GKK(1), GJR(1) |
| GCV | 18 | 0 |
| GJR | 9 | OKV(4), KBB(1), GSK(1), RLC(1), OL3M(1) |
| GKK | 20 | 0 |
| GSK | 16 | OKV(4), GJR(1), RFV(1) |

Individual-level population structuring using a Bayesian approach indicated that the variation among individuals could best be explained if samples were separated into three groups (Figures 4, S1). Again, GCV was the most distinct of the sampled populations. Populations from the Columbia River basin in the USA (GJR and GSK), along with the population from the Klamath River basin (GKK) showed similar group membership in this analysis. The Okanagan Lake populations KBB, LKB, and OL3M were a distinct group, and the remaining Okanagan River Basin populations (OKV, RFV, and RLC) were a mixture between the USA Columbia River populations and the Okanagan Lake populations.

## Discussion

### Overview

To date, two of the four major lineages of freshwater mussels in western Canada and the western contiguous US have been characterized with respect to rangewide genetic divergence and diversity: *Margaritfera falcata* (Mock et al. 2013) and the clade referred to here as *Anodonta nuttalliana* (Chong et al. 2008, Mock et al. 2010). Despite the great distributional overlap between these lineages, divergence and diversity patterns are quite distinct, and are likely shaped by distinct fish host associations and distinct life histories. In both lineages, rangewide diversity was concentrated in northwestern Washington, with lower diversity inland (Mock et al 2013). In *A. nuttalliana*, populations were largely structured by major hydrologic basins (Bonneville, Columbia River, Lahontan), and populations in the Pit and Owens Rivers in northern California were distinct from all other sampled populations according to both nuclear and mitochondrial marker systems (Mock et al 2010). Inbreeding (*FIS*) in *A. nuttalliana* populations was only detectable in two of the 24 populations assessed (Mock et al 2010). In *M. falcata,* by contrast, little clustering was evident among the 48 populations assessed rangewide across much of the same landscape (Mock et al. 2013). High levels of inbreeding were detected in *M. falcata* populations, with *F_IS_* over 0.5 in 33 of the 48 populations, attributable to hermaphroditism (Kat 1983). Here we compare patterns found using similar approaches in *G. angulata* over much of the same landscape, and also examine fine-scale genetic structure within the northernmost populations of *G. angulata* in Canada.

### Interspecific patterns of genetic divergence in western North American freshwater mussel species

We did not find evidence of deep genetic subdivision within *G. angulata.* None of the COI haplotypes differed by more than 4 bp of the 537 bp sequence (0.7%), and there was a continuum of genetic divergence across the Columbia Basin populations (Figure 3). Populations in the Klamath Basin in Oregon did show evidence of isolation, with the majority of samples (31/39) having GonH or GonI mitochondrial haplotypes that were never detected elsewhere (Figure 2). Microsatellite data also showed evidence of moderate divergence in the Klamath (GKK) population (Figure 3). Given that the Klamath Basin is hydrologically distinct from the Columbia River Basin, the degree of genetic similarity (Figure 4) is surprising, although similar patterns were found in *M. falcata* across the Columbia and Klamath Basins as well as other basins in northern California. The Chehalis River population of *G. angulata* in NW Washington, while geographically closer to Columbia River drainages, were strikingly divergent from all other *G. angulata* populations (Figures 3,4) according to nuclear microsatellite markers, and contained a unique haplotype (GonF) at high frequency (3/5 samples) in addition to the ancestral haplotype (GonA) (Figure 2). This contrasts with the pattern found for *M. falcata* (Mock et al. 2013), where the Chehalis River population was not divergent from the Columbia Basin populations. It is possible that anadromous fish host mobility has contributed to divergence patterns in both species, and while we lack the data to test that hypothesis specifically, it does highlight the importance of research into suitable fish hosts for *G. angulata*. Although sculpin (*Cottus* spp.) are believed to be a primary host for *G. angulata,* other fish species may serve as important long-distance dispersers.

### Inbreeding, bottlenecks, and life history across western North American freshwater mussel species

In *G. angulata*, inbreeding estimates were not consistent with high rates of hermaphroditism, but may suggest that low levels do occur, consistent with other observations (van der Schalie 1970). All populations tested had low but positive fixation indices (Table 4) and widespread heterozygote deficiencies (Table 3), suggesting slightly elevated levels of inbreeding. The fixation indices were noted in OKV and KBB, both in the Okanagan River Basin complex. There were no identical genotypes or population bottlenecks detected in the microsatellite dataset in any *G. angulata* populations, although small sample sizes could have limited the power of these analyses. These findings are in stark contrast to findings in *M. falcata*, where multiple identical genotypes were detected within several populations and levels of inbreeding were very high (Mock et al. 2013). The findings in *G. angulata* were also in contrast with *A. nuttalliana*, where inbreeding was generally not detectable, but bottlenecks were detected in a third of the populations tested (Mock et al. 2010). These patterns likely reflect life history differences among species, and may also be influenced by differences in degradation of species-specific habitats and host fish populations.

### Diversity across western North American freshwater mussel species

Mitochondrial COI haplotype richness was higher across *G. angulata* populations (avg. 2.32 haplotypes/22 populations) than in either *M. falcata* (avg. 1.61 haplotypes/18 populations; Mock et al. 2013) or *A. nuttalliana* (avg. 1 haplotype/39 populations; Mock et al. 2010) where samples size was 5 individuals. Microsatellite diversity cannot be compared across species directly, since each species is analyzed with a different set of loci.

### *Diversity within* G. angulata

Across sampled *G. angulata* populations, the most diverse populations in terms of microsatellite allelic richness were those in Oregon (GJR), Vaseux lake in the Okanagan River Basin complex (OKV), the Similkameen River in Washington (GSK), and the Klamath Basin (GKK). The least diverse populations were in the northern portion of Okanagan Lake (KBB) and the Chehalis River (GCV). These results were consistent with haplotype richness (Table 2). This pattern is in contrast with the landscape diversity patterns in *M. falcata* and *A. nuttalliana*, where diversity was concentrated in northwest Washington and the Chehalis River. We had insufficient sample populations in this study to address more regional patterns of diversity in *G. angulata,* but the analysis of additional populations and drainages could provide improved conservation guidance.

### Divergence among Okanagan River Basin populations

Within the Okanagan River Basin complex, our findings suggested a high level of gene flow among most populations. The most isolated population in this complex, KBB, was distinct in the PCoA plot of microsatellite variation (Figure 5), showed evidence of low-level inbreeding (see above) and was the only population in the basin to show 100% self-assignment (Table 5). Other populations in this complex generally clustered together and showed mixed self-assignment. This may suggest a pelagic fish host, in addition to *Cottus* spp. (benthic), since LKB and OL3M are on opposite sides of Okanagan Lake. The KBB population did not, however, show any COI divergence, and only the GonD haplotype was detected, which is the most common haplotype in the Okanagan River Basin complex. These results suggest that (a) the KBB population may have been established by a rare long-distance dispersal event instead of ancient vicariance, and (b) that gene flow within Okanagan Lake may be impeded by the dearth of suitable mussel habitat separating extant RMRM populations at the north and south ends of Okanagan Lake. Apart from a very few mussels noted at one unsampled site in West Kelowna, the northern and southern Okanagan Lake populations are separated by 75 km. The intervening shoreline is mostly steep and rocky, unlike the sandy, gently-sloped beaches that prevail at the north and south ends of the lake, and KBB is a small, spatially isolated population. Microsatellite analysis using STRUCTURE did not support the isolation of KBB from LKB and OL3M, the nearest other populations in Okanagan Lake. The STRUCTURE clustering result does, however, suggest that Okanagan Falls could be an upstream velocity barrier in the Okanagan River Basin complex (Beacham and Withler 2017; Fulton 1969).

### Diversity among Okanagan River Basin populations

Within the Okanagan River Basin complex in British Columbia, there was some signal of increasing microsatellite diversity downstream, consistent with findings in *M. falcata* (Mock et al. 2013) and *A. nuttalliana* (Mock et al. 2010). The Similkameen River population in the U.S. (a major tributary to the Okanogan River in northern Washington) had higher allelic richness than any of the Okanagan River Basin complex populations. LKB and OL3M had intermediate allelic richness, followed by KBB (Table 4). With rarefaction to n=5, allelic richness in RFV and RLC was intermediate between OKV and OL3M. These findings suggest that the strong general downstream diversity patterns seen in *M. falcata* and *A. nuttalliana* may also be present in *G. angulata,* although low numbers of sample populations outside the Okanagan River Basin limited the scale of our inference.

### Biogeographic Implications

Hydrological events impacting drainage patterns in the western United States and Canada have had a profound influence on freshwater fish distributions (e.g., Hershler et al. 2002; Markle 2019; McPhail and Lindsay, 1970, 1986, Minckley et al. 1986), and consequently host availability for native mussels. Markle (2019) presents a review of these geological events, and their implications for the diversity of non-anadromous, true freshwater fishes in Pacific U.S. rivers, notably events leading to the long isolation of the Klamath and Chehalis faunas. In brief, Great Basin volcanism, and more specifically the early Miocene Grande Ronde Basalt Flows (16.8 to 15.5 million years ago) inundated much of the Columbia Basin, including essentially all lands lying between the Klamath and Chehalis watersheds. This would have extirpated the pre-existing lower Columbia River fish fauna, and isolated the Klamath and Chehalis fish and mussel populations. Mountain building, including the Pliocene/Pleistocene uplift of the Klamath Mountains (in the last 5.3 million years) served to further isolate the fish and mussel populations in the Klamath and Chehalis watersheds. The Chehalis fish fauna is said to be “no older than the Pliocene” (2.6 to 5.3 million years ago) (Markle 2019). During this time the Columbia River also re-established a path to the Pacific.

This hydrogeologic history suggests that the ancestral *G. angulata*, represented by haplotype GonA in our data, was perhaps widely distributed among these three watersheds prior to the Grande Ronde Basalt Flows, but over time new, unique haplotypes appeared in each of the three basins: GonF in the Chehalis watershed, GonH in the Klamath Basin, and GonB in the Columbia. The presence of GonB in the Humboldt River (part of the Lahontan Basin) hints at a former connection allowing an exchange of fauna between the Columbia and Lahontan Basins. In *M. falcata,* Mock et al. (2013) found somewhat different patterns: the Klamath and Lahontan populations contained non-overlapping mitochondrial haplotypes, but the Klamath populations shared mitochondrial haplotypes with the Puget Sound region (including the Chehalis River). In *M. falcata*, Neither the Klamath nor Chehalis River populations showed pronounced divergence from other lower Columbia River basin populations with respect to mitochondrial or microsatellite-based metrics. Together these findings suggest that there is little correspondence between patterns of structure between *M. falcata* and *G. angulata*, suggesting that colonization histories, population dynamics, and fish host associations likely contribute to disparate patterns of genetic divergence across a common landscape.

In contrast to the Chehalis, Klamath and Columbia faunas, the Okanagan fish and mussel faunas have very recent origins. The Okanagan Valley had been inundated by the Cordilleran Ice Sheet through the height of the last glaciation. The Okanagan has only been populated (or repopulated) by fish and mussels over the last 10,000 to 15,000 years, via upstream dispersal from the Columbia River and potentially to some extent from the Fraser headwaters (Beacham and Withler 2017). Given the comparative ages and current connectivity, it is not surprising that the Okanagan fauna is little diverged from that of the Columbia River or its many tributaries, as evidenced by both mitochondrial and microsatellite data (Figures 2, 3). Microsatellite allele frequencies among Okanagan Valley populations did provide evidence of population differences (potentially due to population founding events and genetic drift) above and below Okanagan Falls (Figures 1b,5). The falls could impede the upstream (but not downstream) gene flow. A bottleneck is inferred to have affected the KBB fauna (the northernmost population in the Okanagan system). That might reflect relatively recent colonization of the KBB site, or other historical events. As noted earlier, gene flow within Okanagan Lake may be impeded by the dearth of suitable habitat separating extant Rocky Mountain ridged mussel populations at the north and south ends of Okanagan Lake, and may be exacerbated by shoreline development and human activity.

### Conclusions

Effective conservation of freshwater mussel populations requires an understanding of the geographic distribution of genetic diversity and divergence, which can guide restoration programs and help prioritize protections.

Given the unique relationship with host fish species, as well as impacts from human activities, conservation plans need to include those related fish hosts, their mobility and ability to navigate barriers from one suitable habitat to another, and address human impacts. General conservation recommendations include (i) avoiding translocation across major hydrologic basins, (ii) identifying and focusing conservation efforts on fish hosts, and (iii) conducting intensive surveys for discovering additional populations, (iv) collecting and analyzing additional samples from populations represented by under 15 individuals, and (v) monitoring known populations. Some of the samples for this study were collected up to 17 years ago (Table S1), and resurveying and resampling may be particularly important for these populations. We did not detect genetic diversity and inbreeding problems in the populations we studied, with the possible exception of KBB. This presents an opportunity for the establishment of long-term monitoring programs to assure that dramatic population losses do not occur and that genetic diversity does not become eroded.

In Canada’s Okanagan River Basin, translocations across Okanagan Falls should be avoided in order to conserve potential local adaptation. That said, if dramatic population declines are observed, population isolation, genetic drift and loss of genetic diversity may be greater risks than outbreeding depression, and such translocations might be justified (Edmands 2007; Houde 2011). Such decisions depend on other evidence for local adaptation (e.g. contrasting habitats, host fish differences, morphological differences in shells, soft tissues, or glochidia).

## Acknowledgements

This project was undertaken with the financial support of the Government of Canada (Ce project a été réalisé aver l’appui financier du government du Canada), and in-kind support from the British Columbia Ministry of Environment and Climate Change Strategy (Conservation Science Section). For additional financial and field support, we would like to thank the British Columbia Ministry of Forests, Lands, and Natural Resource Operations, and the University of British Columbia – Okanagan Campus. We acknowledge the Confederated Tribes of the Umatilla Indian Reservation, Jayne Brim-Box, Donna Nez, Jeanette Howard, Molly Hallock, Jon Anderson, for providing samples for the U.S. populations of *G. angulata* used in this publication. In addition, we are grateful to Sean MacConnachie (Fisheries and Oceans Canada) for providing feedback during the writing process. We are also grateful to Emilie Blevins (Xerces) for conservation status information.

## Supplementary Materials

Supplementary Materials: https://doi.org/10.26078/fgmq-bm18

Document S1: Microsatellite Development Narrative

Document S2: Raw Microsatellite Sequence Data

Document S3: Screening Scripts

Document S4: Filtered Sequences

Document S5: Primer Design

Document S6: Microsatellite Genotype Data

Figure S1: Structure Harvester Results

Table S1: Sampling Locations

Table S2: Microsatellite Loci and Multiplexing

Table S3: COI Genbank Accessions

## Author Contributions

KEM coordination of molecular analyses, coordinator of genetic analyses, co-lead author. JAW development of microsatellite loci, generation of genetic datasets, co-lead author. SFRB Okanagan fieldwork and sample collection, manuscript co-author.

JHM project design, assistance in funding acquisition, field work coordination, manuscript co-author.

GW project initiation, assistance in funding acquisition, project coordination and field design, manuscript co-author.

IRW assistance in funding acquisition, fieldwork assistance, preparation of figures, interpretation of zoogeography.

---

**Figure S1**. Following the delta-K method of Evanno et al. (2005), implemented in Structure Harvester (Earl & von Holdt 2012), the optimal number for k (the number of clusters of individuals) was 3.

